# Metabolism in the Solar Photon Field A physical framework for photon-assisted modulation of mitochondrial electron-transfer kinetics

**DOI:** 10.64898/2026.08.11.744135

**Authors:** Robert A. E. Fosbury, Roger Seheult, Scott Zimmerman, Glen Jeffery

## Abstract

Within a single human lifetime, the spectral environment has been fundamentally reshaped. Broadband daylight, rich in infrared (IR) photons arising from solar and atmospheric physics, has been replaced in the built environment by narrow, engineered spectra that largely exclude long wavelengths. While modern lighting is optimised for vision, the non-visual photobiology of metabolism may depend on spectral components that are now absent.

When expressed in photon-energy units, the solar spectrum exhibits a broad maximum near 0.75 eV. This range overlaps with the activation and reorganisation energies governing mitochondrial electron-transfer kinetics. Within a Marcus-type framework, IR photons are therefore positioned to modulate rate-limiting metabolic steps by biasing barrier-crossing probabilities rather than supplying chemical energy. These wavelengths also penetrate deeply into tissue in a scattering-dominated regime, forming a diffuse internal photon field capable of interacting with distributed mitochondrial networks.

We propose the term *photometabolism*: a solar-driven, non-photosynthetic modulation of core metabolic processes. A scaling analysis shows that photon interception in this band varies with body mass in parallel with basal metabolic rate, suggesting that ambient sunlight provides sufficient flux to influence metabolic kinetics across the biosphere. These findings have implications for physiology, ecology and the design of indoor environments whose lighting spectra increasingly diverge from their evolutionary context.

## 1. Introduction

Life on Earth evolved under a broadband solar spectrum shaped by stellar physics and atmospheric filtering. Biological science has historically focused on the narrow photic domain of visible wavelengths that drive photosynthesis and vision. The remainder of the solar spectrum that reaches the biosphere has received comparatively little attention in physiology, biochemistry and ecology, despite representing a major component of the natural spectral environment.

Biological science has successfully incorporated temperature, atmospheric composition, gravity and chemistry into its understanding of metabolism. Yet one environmental variable has remained largely absent from this framework: the structured solar photon field within which life evolved.

In *What is Life?* (1944) [81], Schrödinger argued that living systems maintain their highly ordered state by occupying metastable configurations protected by activation barriers against thermal disorder. Although he discussed these barriers primarily in the context of hereditary information, the same physical principle underlies the kinetic stability of metabolism itself.

The present work asks whether these activation barriers, introduced by Schrödinger as the physical basis of biological stability, also provide the point at which metabolism remains coupled to the spectral environment within which life evolved. We therefore explore whether the spectral properties of sunlight may influence electron-transfer kinetics by interacting with the vibrational environment that governs barrier crossing.

The Sun is not simply a broadband radiator. Like other F-, G- and K-type stars, its spectrum exhibits a characteristic enhancement in photon number that closely overlaps both the activation energies inferred for metabolism and the optical window through which biological tissue transmits light most effectively. As discussed below, this enhancement arises from the H⁻ opacity minimum that shapes the continua of Sun-like stars, suggesting possible implications for the spectral environments on habitable planets.

This paper therefore examines, in sequence, the experimental evidence for spectral modulation of metabolism, the structure of the solar photon field, the optical transport properties of biological tissue, the energetics of mitochondrial electron transfer, and their integration within a common physical framework. We do not propose replacing established biochemical pathways, but rather complementing them with a physical perspective in which metabolism is viewed as operating within a structured spectral environment that has accompanied life throughout its evolutionary history.

## 2. Evidence for spectral modulation of metabolism

The hypothesis introduced above is motivated by a broad body of experimental observations that are difficult to reconcile within existing mechanistic frameworks. These studies span diverse species, tissues and experimental paradigms, yet consistently indicate that mitochondrial metabolism responds to illumination over a wide range of longer wavelengths. Before considering possible physical mechanisms, we first summarise the principal experimental evidence that motivates the hypothesis developed in this paper.

In the 1950s, Harman proposed the mitochondrial theory of ageing. The theory argued that ageing arose from oxidative damage to mitochondria, targeting mtDNA and leading to reduced production of ATP and elevated reactive oxygen species (ROS) that drive systemic inflammation. The theory was subsequently refined to propose a self-reinforcing cycle of mitochondrial damage, reduced ATP production and elevated ROS that progressively accelerates cellular ageing [1]. Because ageing is associated with diseases that become increasingly prevalent in older populations, any ability to control this mitochondrial process would offer significant benefits.

Red and near-IR light, approximately 650–900 nm, has been used extensively in animal models of disease and ageing for approximately 20 years to influence mitochondrial regulation of metabolism, ageing and overall health [2–5]. These longer wavelengths, which can penetrate deeply through the body, act directly on mitochondria, although the mechanism of action remains unclear. This light has been thought to influence mitochondrial behaviour via absorption by cytochrome c oxidase in the respiratory chain, thereby improving respiration [5], although more recent studies have argued that the key absorber might be water in the mitochondrial microenvironment because effective IR wavelength ranges overlap with patterns of IR water absorption [6]. The framework developed below may help resolve this issue.

Mitochondria play a broad role in cell signalling and metabolism. They also regulate the pace of ageing. They respond positively to longer-wavelength light in both the near- and mid-IR ranges [7,8], but mitochondrial function is undermined by shorter wavelengths, particularly 420–450 nm [9,10]. Across both vertebrates and invertebrates, longer-wavelength illumination consistently increases mitochondrial membrane potential and ATP production while reducing ROS and inflammation [11–14].

Changes in response to IR are rapid and may be detectable within 3 minutes [15,16], but appear to occur in two stages. Initially, increased ATP production may reflect altered ATP synthase activity, including changes in rotor dynamics. However, at later stages, protein synthesis is initiated, resulting in increased concentrations of mitochondrial proteins, consistent with the production of more respiratory chain elements. Thus, these changes appear to be at the level of the organelle rather than at the level of mtDNA [14]. The changes in response to local light exposure are transmitted across the body, partly via serum cytokine signalling, although this is unlikely to be the only means of transmission [17]. In this way, the effects of IR can become systemic when only limited distal regions of the body are illuminated. A key example of this is the systemic reduction in human blood glucose following IR exposure to relatively small regions of the body. This increases mitochondrial demand for carbohydrates, but the magnitude of the reduction in blood sugars is much greater than that which could occur from the small region exposed to the IR alone [18].

Consistent with Harman’s theory of ageing [1], IR exposure in short-lived animals results in a significant increase in mean lifespan. Maximum lifespan does not change, but the age-related mortality during middle age is markedly reduced. Animals exposed to IR have increased levels of ATP and reduced ROS in middle and old age [11]. They also show improved mobility, cognition and vision during ageing; declines in these functions are significant hallmarks of ageing [16]. These comprehensive improvements reflect the fact that membrane pumps in neurons are among the largest consumers of ATP across the nervous system.

The retina is an ideal model system for ageing because it has the greatest metabolic rate in the body and the highest concentration of mitochondria. Metabolic rate and ageing are linked, with higher metabolic rates having faster ageing patterns [19,20]. Hence, in the primate retina there is a 70% decline in ATP over the lifespan and a 30% loss of central photoreceptors [21,22]. This is reflected in a significant reduction in visual performance as measured psychophysically [23]. IR wavelengths that improve retinal ATP in animals also significantly improve colour vision in older humans. This has been achieved with similar success using IR LEDs at both 670 nm and 850 nm, implying that specific IR wavelengths may not be uniquely important [15,24]. Using specific IR LEDs, single 3-minute exposures have a positive impact on colour vision that lasts approximately 5 days [15]. Similar timescales are found physiologically in insects following IR exposure [25]. Hence, there is a conserved mechanism in this respect. This appears to have switch-like characteristics rather than a simple dose-response relationship to IR light [25]. Different energy levels appear to have similar effects. Consequently, IR wavelength and exposure duration may not be critical variables if there is enough available energy to trigger the response [7]. Studies using insects indicate that the exposure required to trigger this switch can be as short as just over 1 minute, and in humans the energy levels may be as low as around 1 mW/cm² [25,26]. However, the time of day may be important. Selective IR exposure appears to have the greatest impact early in the day, when ATP production is dynamic. This pattern is conserved between flies and humans. Exposures later in the day have less impact [15,27].

Despite consistent positive data, there is growing evidence that light from the restricted spectral range of IR LEDs is suboptimal. When people are exposed to incandescent lighting, with its broad, continuous spectrum extending throughout the IR range, similar to sunlight, the impact on visual ability is more profound and lasts much longer after the light is removed than after IR LED exposure [15,28]. This point is significant because it highlights the value of sunlight with its continuous IR spectrum and the potential need to reproduce or approximate this spectrum when trying to manipulate mitochondrial function optically.

Metabolic decline is a common feature of many diseases, irrespective of whether those diseases have a primary mitochondrial basis. Mitochondrial dysfunction in brainstem and dopaminergic pathways has been implicated in Parkinson’s disease pathophysiology, making these pathways a target of much IR research. Experimental models of Parkinson’s disease similarly show significant improvements following either direct brainstem or peripheral IR exposure, consistent with systemic signalling [29].

Ageing results in mitochondrially mediated apoptosis. Here, the mitochondrial membrane becomes permeabilised and leaks cytochrome c, activating caspases and leading to cell destruction. IR exposure that significantly improves membrane potential is associated with significant reductions in cell death in both normal retinal ageing and experimentally induced Parkinson’s disease [30,31]. Hence, from an ageing perspective, IR-induced improvements in mitochondrial function are associated with better lifespan-related outcomes, improved mobility and metabolism, reduced inflammation, and a slower pace of age-related cell death. The magnitude of these improvements is often statistically significant. One possible reason is the background lighting conditions in which these studies take place. Laboratory animals and human subjects are often housed or studied in modern lighting environments largely devoid of IR, with illumination often provided by fluorescent strip lighting or standard LEDs whose emission spectra rarely extend significantly beyond 650 nm. Thus, mitochondrial light-responsive pathways may be relatively under-stimulated in such environments. A key experiment yet to be undertaken would be a direct comparison between sunlight and modern LEDs, while holding all the other conditions constant.

An important and independently established pathway linking IR radiation to mitochondrial function involves nitric oxide (NO). Photodissociation of NO from cytochrome c oxidase has been shown to relieve inhibition of mitochondrial respiration, thereby increasing electron transport and ATP production [32]. In parallel, longer-wavelength irradiation can mobilise NO from S-nitrosothiol reservoirs, leading to vasodilation and enhanced tissue oxygenation, which is critical for mitochondrial function. These processes provide a direct physiological link between light exposure, mitochondrial activity, and systemic metabolic regulation, and they are supported by a substantial biochemical and clinical literature.

Although nitric oxide pathways account for several observed effects, they do not, by themselves, explain key features of the experimental data. They do not readily account for the broad spectral sensitivity across the near- and mid-IR, the comparable efficacy of different wavelengths, or the persistence of responses following brief exposures. These observations suggest that NO-mediated processes form part of a wider framework in which IR radiation interacts with mitochondrial function through more general physical mechanisms.

Collectively, these observations point towards a conserved mitochondrial response to longer-wavelength illumination that extends across species, tissues and physiological contexts. They also reveal several features that are not readily explained by existing mechanistic models, including broad spectral sensitivity, comparable responses across different wavelengths, systemic effects following local exposure, and biological responses that persist long after illumination has ceased. These recurring characteristics motivate the search for a more general physical framework capable of linking the diverse experimental observations reviewed above.

## 3. Clinical and epidemiological evidence

Independent testing of the proposed framework at organismal and population scales is important. If IR-rich sunlight modulates mitochondrial electron transfer and redox balance, corresponding effects should be detectable not only in physical models and cellular systems but also in physiology, clinical outcomes, and epidemiological patterns. The evidence reviewed below is presented as convergent evidence across biological scales, although it does not by itself establish the mechanism.

Mitochondrial function is central to the pathophysiology of many common chronic conditions, including heart failure [33], diabetes mellitus [34], obesity [35], neurodegenerative disorders [36,37], cancer [38], infectious diseases [39,40], and ageing itself [41]. In these conditions, impaired electron-transfer efficiency and excess ROS production are recurrent features, linking mitochondrial dysfunction to both metabolic decline and tissue injury. These processes therefore provide an important context within which to examine the physiological consequences of the light-dependent modulation described in the previous section. If IR photons within the solar spectrum can modulate electron transport to improve mitochondrial efficiency and reduce ROS leakage, as suggested by the physical framework developed below, then corresponding signatures should be observable in clinical, epidemiological, and experimental data.

Experimental studies demonstrate that exposure to IR accelerates whole-body metabolic rate, as reflected by increased end-tidal CO₂ and reductions in blood glucose [18]. These observations are consistent with enhanced mitochondrial electron flux rather than with photochemical or thermal effects. At the population level, large observational cohorts in Europe report associations between greater sunlight exposure and reduced all-cause, cardiovascular, and cancer mortality [42,43]. In one cohort of approximately 10,000 participants, greater hourly sunlight exposure was associated with improved insulin sensitivity and lower triglyceride levels [44]. While this cannot establish causality, it suggests a widespread systemic physiological influence of sunlight.

IR-rich sunlight exposure has also been repeatedly linked to infectious-disease susceptibility and outcomes among infected individuals [45]. Seasonal influenza often peaks shortly after the winter solstice in temperate regions, while equatorial regions exhibit little seasonal variation. The unusually early onset of the 2009 H1N1 pandemic enabled partial separation of the effects of temperature and solar radiation; analyses integrating solar radiation measurements with influenza surveillance data concluded that sunlight exposure was associated with a strong protective effect [46]. Parallel patterns have been reported for COVID-19, with incidence and mortality correlating more strongly with latitude and ultraviolet radiation than with temperature or humidity, and with effects that appear independent of vitamin D production [47–49].

Interventional studies provide further support. Controlled trials using IR illumination have reported improved clinical outcomes, including reduced length of hospital stay and enhanced functional recovery [50]. Randomised, placebo-controlled and blinded studies in intensive-care settings using combinations of red and near-IR light have reported reductions in length of stay and improved muscle strength at discharge [51]. Observational studies similarly report shorter hospital stays and better outcomes in rooms with greater access to daylight [52–54].

Importantly, several studies indicate that broadband IR exposure, rather than monochromatic or narrow-band illumination, produces more robust physiological effects [28]. In workplace and home settings, polychromatic light sources containing a continuous IR component have been associated with improved colour perception, mood, alertness and cardiometabolic markers, particularly during winter months when natural sunlight exposure is limited [55]. These findings align with the physical argument developed here: namely, that modulation of mitochondrial function depends on coupling to a distributed vibrational environment, rather than excitation at a single wavelength.

Taken together, these epidemiological associations, physiological measurements and clinical interventions indicate that the experimental responses described in Section 2 extend beyond isolated laboratory systems to whole organisms and human populations. Although they do not establish the underlying mechanism, they reinforce the need for a physical explanation capable of linking observations across these multiple biological scales. The following sections therefore examine whether the spectral structure of sunlight itself provides that missing framework.

## 4. Solar spectrum and the evolutionary photic environment

The experimental and clinical observations summarised above reveal a conserved, broadband and persistent influence of long-wavelength light on mitochondrial function across diverse organisms and tissues. These observations are not fully explained by classical photochemistry, specific chromophore absorption or bulk thermal effects, suggesting that a more general physical process may be involved. The common denominator underlying these diverse responses is the mitochondrial electron transport chain (ETC), whose activity depends critically on electron-transfer reactions occurring within a hydrated protein–membrane environment.

We therefore develop a physical framework, termed *photometabolism*, in which naturally available solar photons modulate metabolic electron-transfer kinetics without contributing directly to metabolic free energy. The following sections examine this physical environment in progressively greater detail, beginning with the spectral structure of sunlight, continuing through its transport within living tissue, and culminating in the electron-transfer mechanisms through which weak photonic perturbations may influence mitochondrial function.

### 4.1 The solar photon-energy distribution

When expressed in conventional radiometric units (W m⁻² nm⁻¹), the solar spectrum appears to peak within the visible range. Many biological processes depend on the number of photons available at a given energy rather than on radiative power alone. Recasting the spectrum as photon flux per unit energy therefore provides a more appropriate representation for processes that involve molecular excitation and electron transfer.

For processes involving individual photons, it is convenient to transform from spectral irradiance, *E _λ_* (*λ*) [W m⁻² nm⁻¹], into photon flux per unit energy, *Φ_E_*(*E*) [photons s⁻¹ m⁻² eV⁻¹]. This transformation is obtained by applying the appropriate Jacobian when changing variables from wavelength to photon energy:

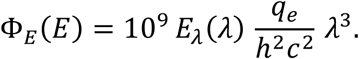

With the spectral data tabulated in 1 nm wavelength bins, this expression accounts for (i) conversion from per nanometre to per metre, (ii) division by the photon energy *E_ph_ = hc/λ*, (iii) transformation from wavelength to energy using the chain rule [56], and (iv) conversion from joules to electron volts through *E_eV_ = E_J_/q_e_*, where *h*, *c,* and *q_e_* are the Planck constant, the speed of light and the elementary charge, respectively. The transformation preserves the radiometric information but redistributes it into equal photon-energy intervals.

Because of the *λ³* Jacobian weighting, a blackbody at the Sun’s effective temperature (*T_eff_* ≈ 5772 K) exhibits a broad maximum near 0.75 eV (approximately 1650 nm) rather than within the visible spectrum (Fig. 1). Although the photon-flux maximum lies slightly beyond the conventional 700–1300 nm tissue window, it adjoins the spectral region over which tissue transmission remains exceptionally high, allowing substantial photon penetration into living tissue. Similar representations are routinely employed in photovoltaic research and are particularly appropriate when considering quantum processes in biology.

**Figure 1.**
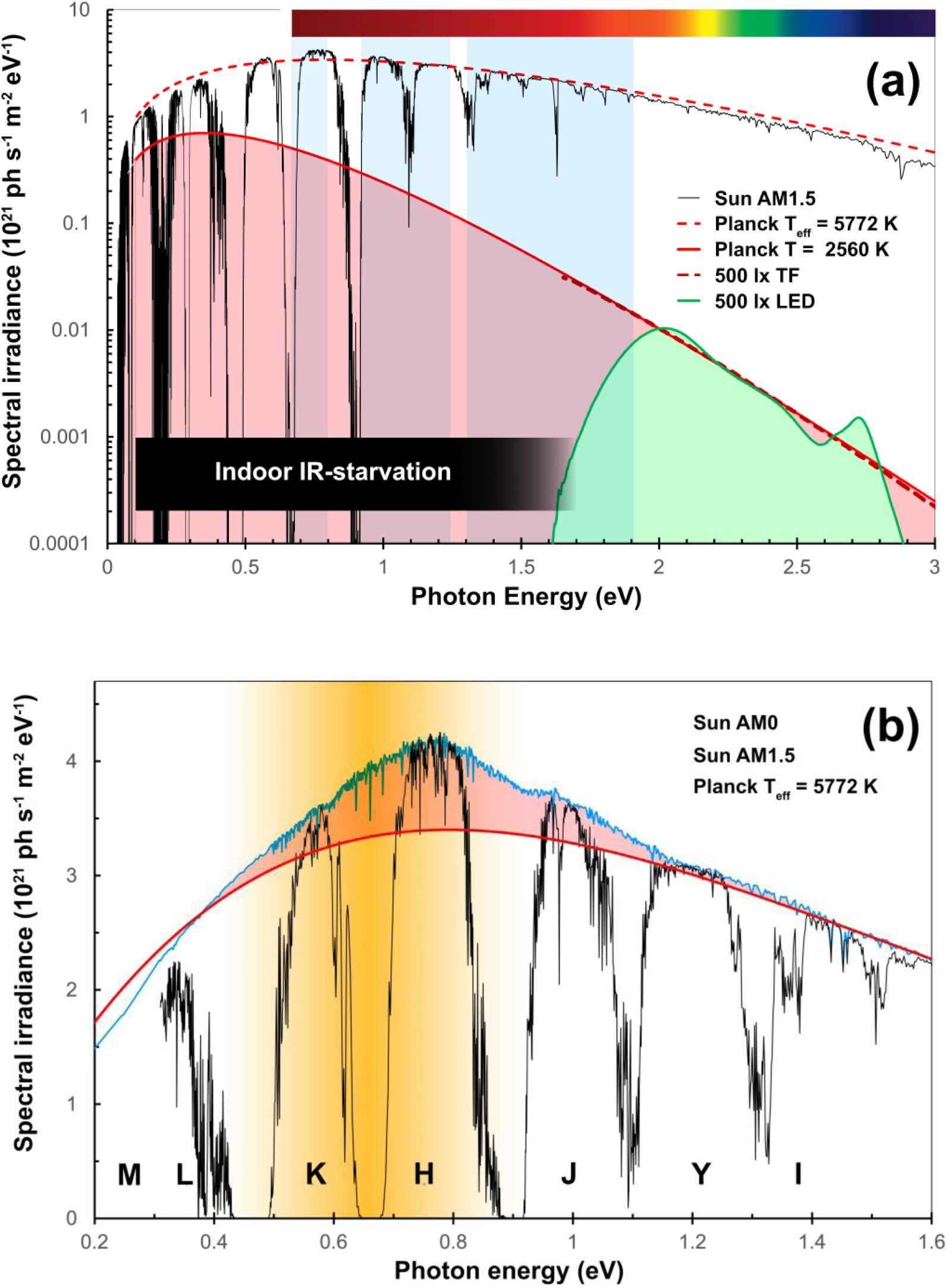
The terrestrial solar photon environment expressed in photon-energy units and its correspondence with metabolic energetics. **(a)** Solar photon flux at the Earth’s surface expressed as photon flux per unit energy (photons s⁻¹ m⁻² eV⁻¹) on a logarithmic ordinate. The terrestrial AM1.5 spectrum (black) exhibits a broad maximum near 0.7–0.8 eV. Atmospheric attenuation is relatively modest across the visible range, whereas water-vapour absorption produces structured bands at lower photon energies and longer wavelengths. Also shown are spectra representative of indoor lighting: a white LED source (green shading) and a tungsten-filament source (pink shading), each scaled to a photopic illuminance of 500 lx. Dashed curves show Planck distributions at 5772 K, the solar effective temperature, and 2560 K, representative of a tungsten filament. Blue shading indicates the biological tissue transmission window (BTTW). The comparison highlights the strong depletion of long-wavelength photon flux under typical indoor lighting conditions relative to sunlight, despite similar visual brightness. **(b)** The same solar photon-flux distribution shown on a linear ordinate to emphasise the broad maximum near 0.7–0.8 eV and its physical origin. The astronomical IR transmission windows are labelled I–M. The red curve is a Planck distribution at the solar effective temperature of 5772 K, normalised to reproduce the total bolometric solar output, while the blue and black curves show the solar spectrum above the atmosphere (AM0) and at the Earth’s surface (AM1.5), respectively. Pink shading highlights the enhancement of the observed solar spectrum above the bolometrically normalised Planck distribution. This enhancement arises from the minimum in H⁻ continuum opacity near the photodetachment threshold at 0.754 eV, which allows radiation to emerge from deeper, hotter photospheric layers. The orange band indicates the characteristic range of activation energies observed across diverse taxa (mean approximately 0.66 eV [80]). Their broad correspondence with the solar photon-flux maximum illustrates the convergence between the terrestrial spectral environment and the energetic scale of metabolism that motivates the photometabolism hypothesis.

Superimposed on this Planckian distribution is a distinctive feature arising from the *H⁻* ion, the dominant continuum opacity source in the atmospheres of the Sun and other F–K stars. Near its bound–free threshold (0.754 eV), the opacity reaches a broad minimum [57], allowing radiation to emerge from deeper, hotter photospheric layers. The resulting enhancement of the emergent photon flux—approximately 20–25% above the corresponding Planck continuum— produces the broad plateau centred near 0.75 eV shown in Figs. 1a and b. Consequently, the dominant photon energies reaching Earth’s surface occupy the same sub-electron-volt regime as the activation barriers inferred for metabolism. This correspondence motivates the hypothesis, explored below, that naturally available solar photons can influence metabolic kinetics through interactions with the vibrational environment governing mitochondrial electron transfer.

### 4.2 Atmospheric filtering and photon transport in tissue

Solar photons are filtered first by the atmosphere and then by biological tissue. Atmospheric transmission in the IR is shaped by rotational–vibrational absorption bands of water vapour, producing the familiar astronomical windows. Within tissue, absorption is smoother and red-shifted, defining a biological tissue transmission window (BTTW) extending approximately from 0.7 to 1.3 µm with some transmittance up to 1.9 µm [58].

Fig. 2 compares the wavelength dependence of diffuse transmittance (DT) and diffuse reflectance (DR) of human tissue with atmospheric transmission. It also includes a typical example of the transmission of currently available—and widely employed—low-emissivity window glass. The comparison of these data with the spectral irradiances of sunlight, a white LED, and a hot-filament lamp in Fig. 1a clearly illustrates the problem of IR starvation in the built environment. Imaging representations of the marked difference in behaviour of visible and near-IR light are shown in Fig. 3 (adapted from [24]).

**Figure 2.**
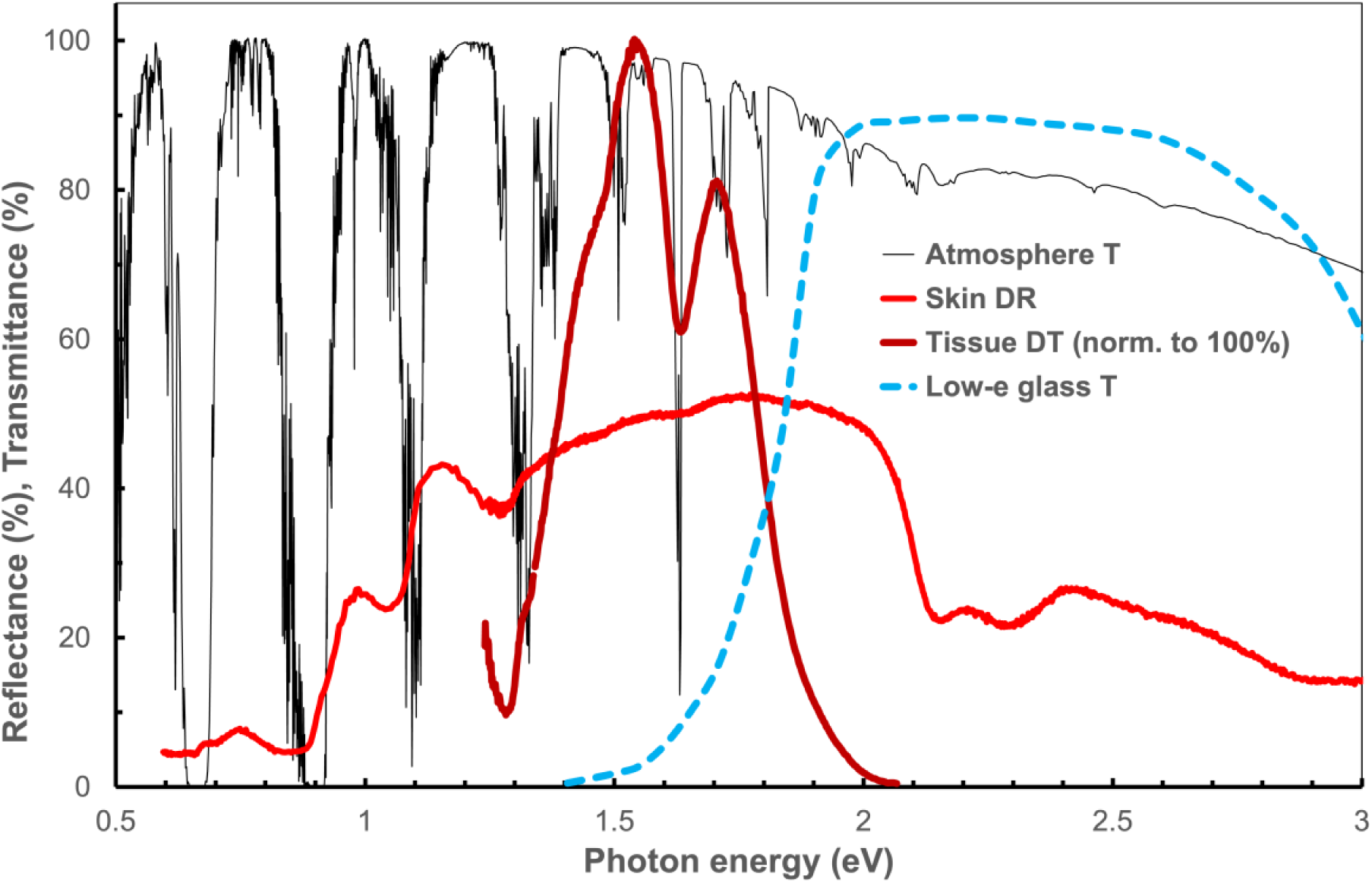
Comparison of spectral transmission through the Earth’s atmosphere (black) and a typical low-emissivity (low-E) window glass (blue dashed), together with optical measurements of human tissue. The red curve shows diffuse reflectance (DR) from human skin, arising from multiply scattered photons that have penetrated and re-emerged from the tissue. The dark red curve shows the corresponding diffuse transmittance (DT), representing light that has undergone multiple scattering events within the tissue before exiting; this curve is normalised to a peak of 100%. Together, these data illustrate the scattering-dominated optical regime of biological tissue and its correspondence with atmospheric transmission windows, as well as the strong attenuation of IR radiation by modern architectural glass.

**Figure 3.**
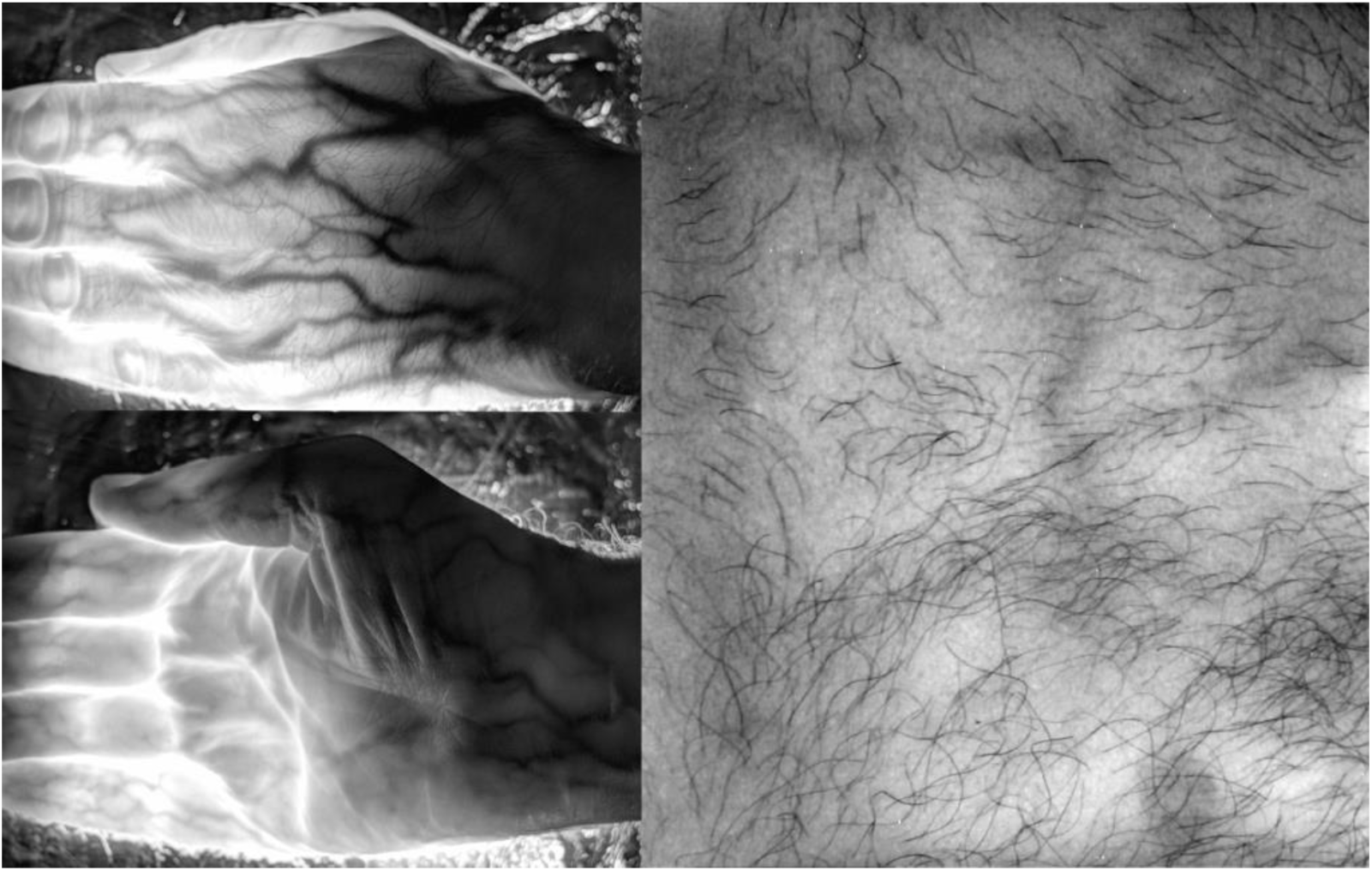
Direct visual evidence for deep penetration of near-IR light (approximately 600–1000 nm) through human tissue. Left: transmission images of the human hand at approximately 850 nm (after [24]). Right: reprocessed transmission image of the human thorax (adapted from [24]), showing substantial passage of sunlight through an estimated thoracic tissue depth of approximately 30 cm. In both cases, high-contrast internal features - including subcutaneous veins and fine hair structure - are clearly resolved, indicating that the detected signal is dominated by multiply scattered, internally transmitted light rather than surface reflection or ambient stray illumination. These observations provide macroscopic confirmation of the effective optical transparency of living tissue within the near-IR BTTW, consistent with deep photon penetration and volumetric light transport.

In this regime, scattering dominates over absorption. Photons undergo multiple deflections, increasing their residence time and effective path length before absorption. Tissue behaves as a low-Q optical cavity in which photons encounter structures with varying degrees of spatial regularity, undergo multiple randomising deflections, and persist for extended residence times — analogous to sound reverberating within a cathedral before being gradually absorbed. This produces a diffuse internal photon field in which even weak absorbers can interact efficiently. The result is a broadband, spatially distributed photon field capable of coupling weakly but collectively to intracellular structures. This transport regime does not favour narrow action spectra. Instead, it supports weak coupling to the collective vibrational modes of water, lipids and proteins that define the intracellular environment and govern electron-transfer kinetics.

Beyond the BTTW, absorption by liquid water rises substantially and photon penetration depths decrease. However, the longer atmospheric windows H, K, L, M, and beyond [59,60] still deliver substantial solar photon flux to the skin surface. Under these conditions, energy deposition becomes concentrated within superficial tissues, where the high local photon flux may influence microvascular dynamics, structural water organisation, connective tissue remodelling and redox balance through pathways distinct from those governing deep mitochondrial coupling.

### 4.3 Spectral divergence in the modern built environment

The modern built environment represents a substantial departure from the spectral conditions under which biological systems evolved. Daylight provides a continuous photon distribution extending from the visible into the near- and mid-IR, with a large fraction of total photon flux residing at wavelengths longer than 700 nm. In contrast, standard white LEDs and fluorescent sources exhibit sharply truncated spectra, with emission typically falling to negligible levels beyond approximately 650–700 nm [58]. This spectral truncation greatly reduces the population of sub-electron-volt photons centred near 0.75 eV (Fig. 1).

This deficit is further compounded by architectural materials. Low-emissivity window coatings, now widely used for thermal efficiency, are designed to reject IR radiation, significantly reducing transmission above approximately 800–1000 nm (Fig. 2). As a result, indoor environments can exhibit reductions of IR photon flux by an order of magnitude or more relative to outdoor daylight, even when visible illumination levels appear comparable.

Taken together, artificial lighting and glazing systems selectively exclude those wavelengths that coincide with metabolic activation barriers and, within the biological tissue transmission window, penetrate living tissue most effectively. The built environment therefore imposes a systematic spectral shift that may decouple metabolic regulation from its evolutionary photic context.

## 5. Mechanisms and scaling

The preceding sections have established that the terrestrial biosphere evolved within a highly structured solar photon environment and that modern lighting substantially modifies that environment. We now consider how such spectral differences might influence metabolism. Our central hypothesis is that near-IR photons do not drive mitochondrial photochemistry directly, but instead perturb the activation barriers governing electron transfer. Small changes in these near-threshold kinetic barriers may then be amplified through physiological signalling and ultimately reflected across the metabolic scaling relationships that characterise life over many orders of magnitude.

### 5.1 Electron-transfer kinetics and barrier modulation

The proposed mechanism is hierarchical in scale. At its foundation lies electron-transfer kinetics, in which reaction rates are governed by coupling between electronic states and nuclear reorganisation. Near-IR radiation is proposed to interact with this system indirectly through excitation of collective vibrational modes of water, lipids and proteins that define the reorganisation coordinate. Subsequent physiological processes, including nitric oxide signalling and perfusion changes, then act as downstream amplifiers linking local modulation of electron transfer to systemic metabolic responses.

At the cellular level, absorption of near-IR radiation in biological tissue is dominated by broad vibrational overtones of water and lipid acyl chains arising from hydrogen-bonded and collective molecular modes [58,62]. These collective solvent and environmental motions form the dynamical background of the intracellular medium and determine the activation barriers governing metabolic processes. Such barriers regulate reaction rates, preventing uncontrolled chemistry while allowing metabolic pathways to operate at physiologically useful speeds.

Metabolic energy is supplied through the mitochondrial ETC, where electrons move sequentially between redox centres embedded in a hydrated protein–membrane matrix. Although the overall flow is thermodynamically downhill, several steps are rate-limited by activation barriers that regulate both metabolic flux and the balance between efficient transfer and ROS leakage.

The proposed mechanism is illustrated schematically in Fig. 4a. Electrons move sequentially through a chain of redox centres embedded within a hydrated protein–membrane environment. Rather than directly promoting electronic excitation, near-IR photons are proposed to perturb the collective nuclear motions of this environment, subtly altering the probability of crossing the activation barriers that regulate electron transfer.

**Figure 4.**
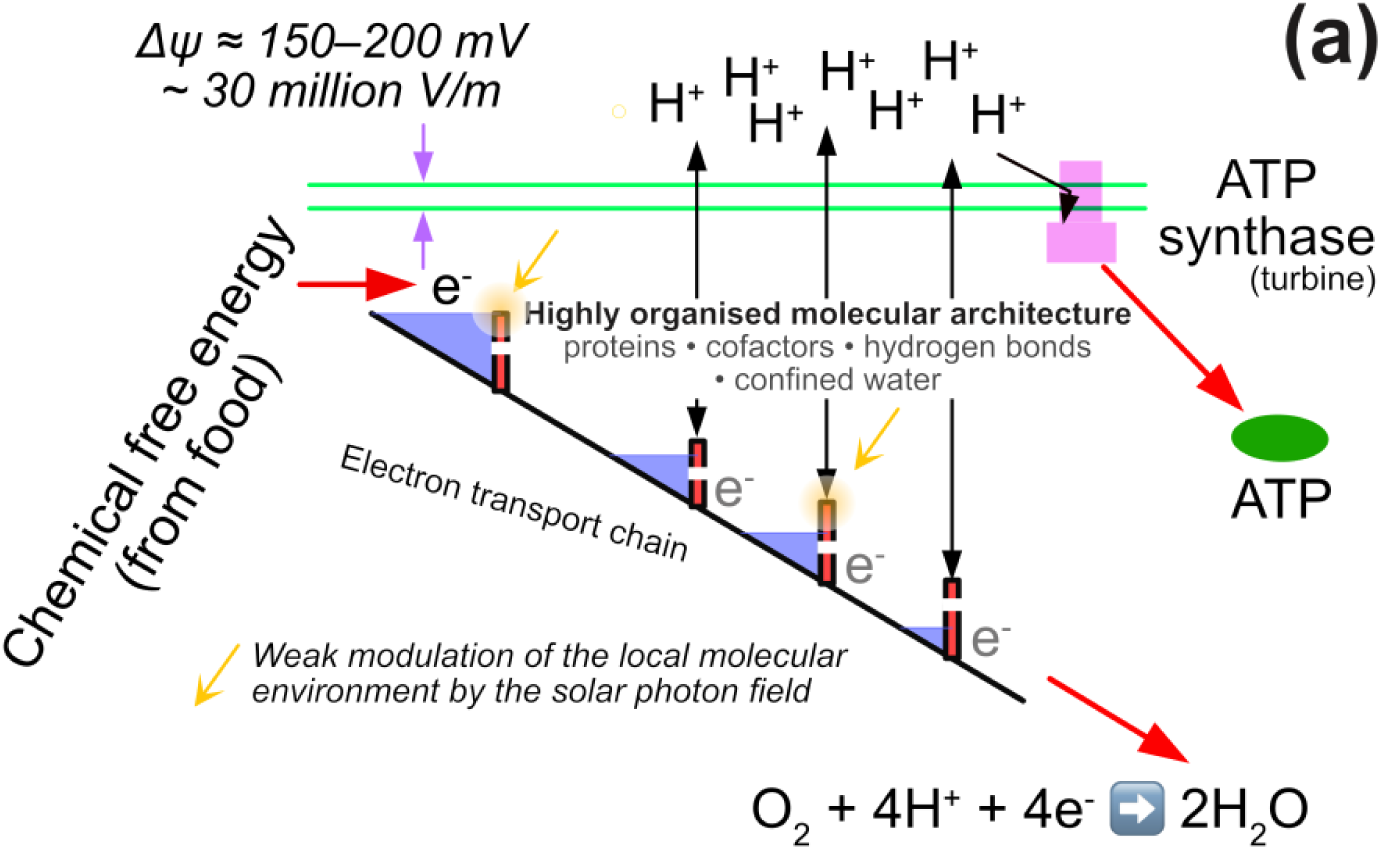
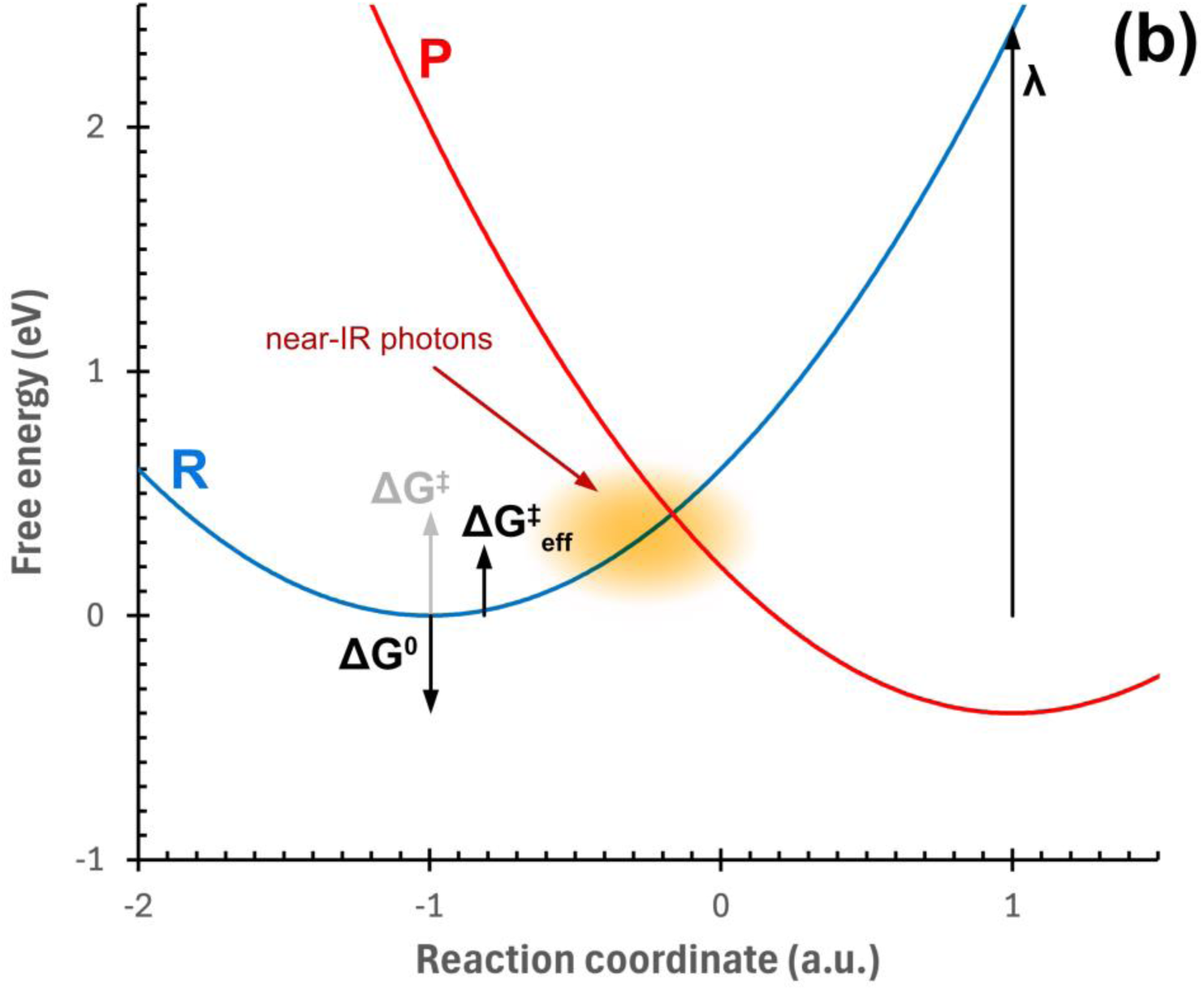
The electron transport chain and Marcus kinetics. **(a)** Conceptual representation of mitochondrial energy conversion and the proposed site of photometabolic modulation. Chemical free energy derived from food drives electron transport through a sequence of Marcus-type activation barriers, generating the proton motive force (Δψ) across the inner mitochondrial membrane that powers ATP synthase. The activation barriers are embedded within a highly organised local molecular architecture comprising proteins, cofactors, hydrogen-bond networks, and confined water. The present hypothesis proposes that naturally available solar photons do not directly contribute metabolic free energy but instead weakly modulate this local molecular environment, thereby influencing electron-transfer kinetics while leaving the underlying biochemical pathways unchanged. The figure is schematic and is intended to illustrate energetic principles rather than the detailed molecular structure of the respiratory complexes. **(b)** Schematic Marcus free-energy surfaces for electron transfer showing reactant (R) and product (P) states, with the reorganisation energy λ, reaction free energy ΔG⁰, and activation barrier ΔG‡ indicated. The shaded region marks the configuration space near the crossing point where electron transfer occurs. Coupling of near-IR to low-frequency vibrational modes is proposed to enhance fluctuations along the reaction coordinate, increasing access to this region and reducing the effective barrier (ΔG‡ → ΔG‡_eff_) without necessarily modifying the underlying free-energy surfaces.

This coupling between environmental fluctuations and reaction rate is formalised in Marcus theory [63, 64], developed for condensed-phase and biological systems by Newton and Sutin [65], and extended to include quantum vibrational modes by Jortner [66], whose formulation incorporates coupling to high-frequency vibrational modes. A comprehensive review, including application to biology, is presented by Marcus and Sutin [67]. Within this framework, the activation barrier for an electron-transfer step is given by ΔG‡ = (λ + ΔG⁰)² / (4λ), where λ is the reorganisation energy and ΔG⁰ the reaction free energy. In the presence of a weak photonic perturbation, the effective barrier may be written ΔG‡(eff) ≈ ΔG‡ − αEγ, where Eγ is the photon energy and α ≪ 1 represents an effective coupling to the reaction coordinate. This phenomenological expression is intended only to illustrate the direction of barrier modulation rather than provide a formal derivation of electron-transfer kinetics. It does not imply direct photon-driven electron transfer, but captures the idea that photon absorption or subsequent vibrational relaxation can shift the system along its reorganisation coordinate, increasing the probability of barrier crossing. In systems operating near threshold (ΔG‡ ∼ 0.5–0.8 eV), even a small reduction in barrier height can produce a measurable increase in electron flux.

The corresponding Marcus free-energy surfaces are shown schematically in Fig. 4b. Electron transfer occurs only when fluctuations of the surrounding nuclear environment transiently bring donor and acceptor states into near resonance, allowing the system to sample the narrow crossing region between the free-energy surfaces. The reaction rate is therefore governed by how frequently this region is accessed rather than by the thermodynamic driving force alone.

Near-IR photons, by coupling to low-frequency collective modes of water and the surrounding biomolecular environment, are plausible candidates for such a perturbation, enhancing access to the crossing region rather than driving discrete electronic transitions.

In this context, electron transfer may be viewed as a sequence of kinetically gated transitions between neighbouring energy states rather than abrupt, high-energy jumps. A useful analogy is that of a canal lock system, in which water levels are adjusted incrementally to allow controlled passage between sections, avoiding large, uncontrolled drops. In biological electron transport, reorganisation of the surrounding protein and hydration environment serves a similar role, aligning energy levels so that electrons move stepwise across modest barriers. Operation near kinetic thresholds minimises the lifetime of poorly coupled or high-energy intermediates, reducing the probability of off-pathway electron leakage that gives rise to ROS. Within this framework, small perturbations — including those induced by red and near-IR photons — may bias these transitions by enhancing flux while maintaining controlled, low-damage operation.

The principle that environmental nuclear motions shape electronic kinetics is not unique to biology. In condensed-matter systems, electron mobility in crystals is influenced by coupling to lattice phonons, while in molecular solutions charge transfer is modulated by solvent polarisation modes. In each case, the surrounding vibrational environment governs the probability of barrier crossing without altering reaction stoichiometry. The mitochondrial matrix and inner membrane form an analogous condensed-phase environment dominated by collective modes of water, lipid, and protein scaffolds.

Within this framework, near-IR photons act not as direct chemical reagents but as weak perturbations of the fluctuating molecular environment that governs mitochondrial electron transfer. This hypothesis provides a physically consistent mechanism linking the structured solar photon field described in Section 4 to the experimentally observed metabolic responses reviewed in Sections 2 and 3. The physiological consequences of such subtle kinetic modulation are considered in the following subsection.

### 5.2 Biological amplification across timescales

If IR photons act by subtly perturbing electron-transfer kinetics, the resulting changes need not remain confined to individual redox reactions. Mitochondrial metabolism occupies a central position within a hierarchy of regulatory pathways, allowing small changes in electron-transfer flux to propagate through successive levels of physiological organisation. The experimentally observed responses to brief IR exposure are consistent with such amplification, extending from rapid metabolic adjustments to longer-term changes in cellular function and tissue repair.

The reported physiological responses span a hierarchy of timescales. Rapid responses occurring within minutes are consistent with immediate modulation of electron-transfer flux and ATP synthesis [10,16]. Intermediate responses over hours involve signalling pathways, including nitric oxide (NO) release, vascular responses and cytokine-mediated communication. Longer-lived effects extending over days are associated with protein synthesis, mitochondrial biogenesis and tissue repair [14,15]. Within the present framework, a weak perturbation of electron-transfer kinetics therefore acts not as an isolated event but as the trigger for a cascade of biological amplification.

One important amplification pathway involves nitric oxide. Red and near-IR irradiation promotes NO release from both cytochrome c oxidase and S-nitrosothiol reservoirs, relieving NO-mediated inhibition of mitochondrial respiration while simultaneously inducing vasodilation and improving tissue perfusion [68–73]. In this way, local modulation of mitochondrial electron transfer may be coupled directly to oxygen delivery, providing a mechanism by which relatively small intracellular changes are amplified at the tissue level.

Emerging evidence further suggests that longer-wavelength IR interactions with collective vibrational modes of water and proteins may provide an additional indirect route for influencing these near-threshold processes [74].

These physiological amplification pathways are distinct from, but complementary to, the Marcus-type kinetic modulation proposed in Section 5.1. Together they provide a plausible explanation for one of the most striking features of the experimental literature: that brief periods of IR exposure can initiate biological responses persisting for hours, days or even longer. Within this hierarchical framework, subtle modulation of mitochondrial electron-transfer kinetics is progressively amplified through metabolic regulation, cellular signalling and tissue-level physiology.

### 5.3 Metabolic scaling and the solar photon environment

If the proposed mechanism contributes to metabolism across the biosphere, it should remain physically applicable over the enormous range of organism sizes represented in nature. Metabolic scaling theory provides an opportunity to examine this possibility. Across organisms, basal metabolic rate follows a near-universal three-quarter power relationship with body mass (*M* ^3/4^), reflecting fundamental constraints on resource distribution and energy use [75,76]. When the solar photon spectral flux is integrated across a 1 eV-wide band spanning 0.31–1.31 eV, it encompasses the broad maximum near 0.75 eV. This range corresponds to the domain in which the solar photon-flux distribution, the characteristic activation energies observed across the biosphere, and mitochondrial electron-transfer barriers overlap.

The quantitative comparison is illustrated in Fig. 5 using three representative organisms for which experimental evidence exists for IR influences on metabolism: Drosophila melanogaster, the mouse and the human. Basal metabolic rates were taken from the literature [77,78]. The available solar power was estimated from the ASTM G173-03 AM1.5 Global Tilt spectrum [79] by integrating over the photon-energy interval 0.31–1.31 eV (947–4000 nm), encompassing the broad maximum of the solar photon-flux distribution near 1650 nm. Multiplying this irradiance (∼280 W m⁻²) by the projected intercept area of each organism provides a first-order estimate of the incident solar power available for interaction. Projected areas, estimated for representative postures at a solar altitude of approximately 42°, are 2 × 10⁻⁶, 3 × 10⁻³ and 0.7 m² for Drosophila, mouse and human respectively. Projected intercept area, rather than total surface area, is the relevant geometric quantity here because it determines the number of incident solar photons available to the organism.

**Figure 5.**
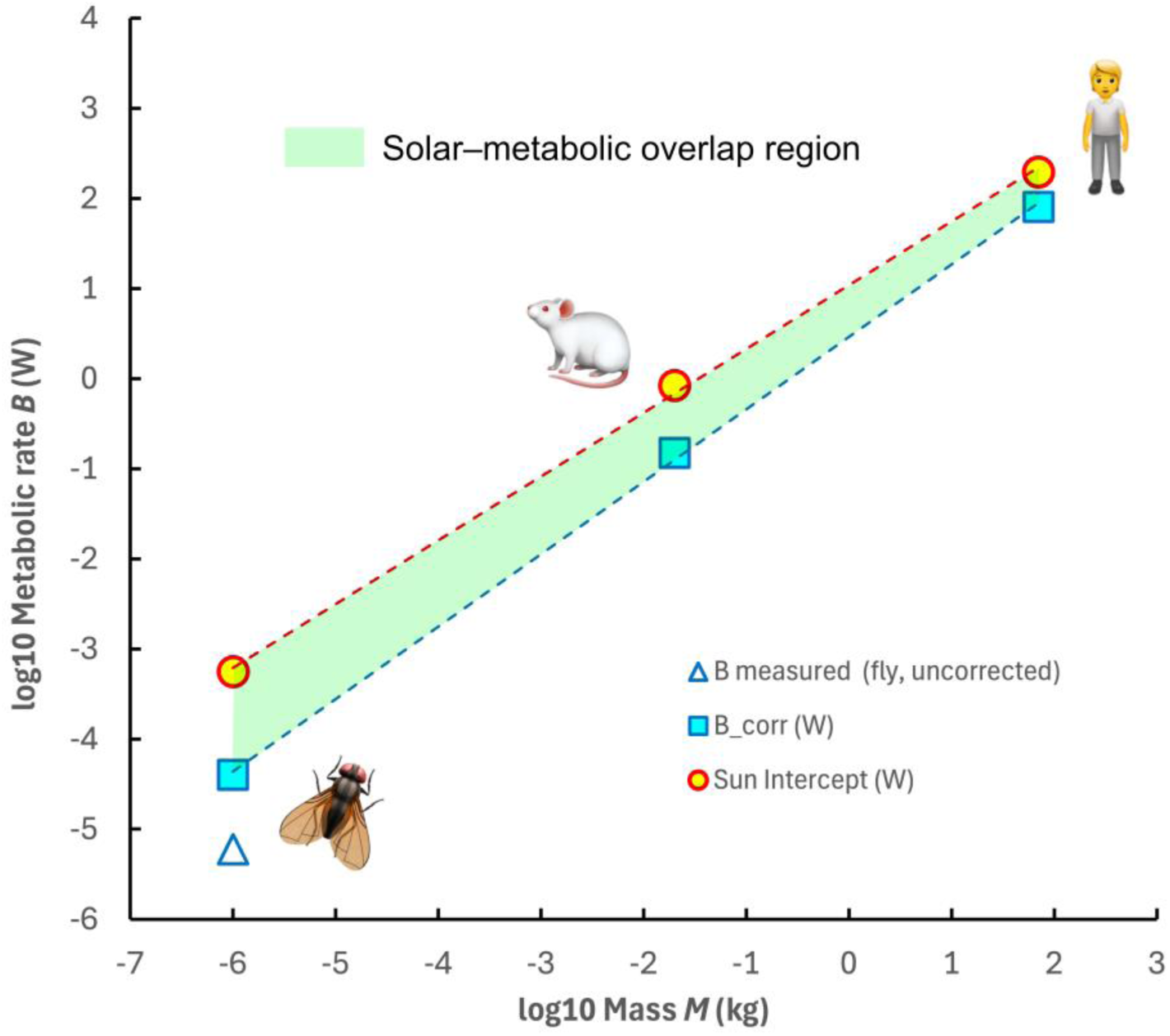
Comparison of basal metabolic rate and available incident solar photon power across body mass. Blue squares show basal metabolic rate following the metabolic theory of ecology (MTE) scaling; the open blue triangle indicates the measured Drosophila value prior to temperature correction, and the filled square its Arrhenius-corrected value. Orange circles show the solar photon power intercepted by the projected body cross-section under AM1.5 illumination at a solar altitude of approximately 42°, integrated over a 1 eV-wide photon-energy band spanning 0.31–1.31 eV and encompassing the broad maximum near 0.75 eV. (Fig. 1). Across eight orders of magnitude in body mass, available incident photon power scales approximately in parallel with metabolic rate and is of the same order as, and for larger organisms may exceed, basal metabolic power. This comparison concerns incident environmental photon power, not metabolically harvested energy, and it is not a claim that solar radiation supplies metabolic demand. Rather, the correspondence supports the possibility that ambient solar photons may influence mitochondrial electron-transfer kinetics through weak but persistent coupling to intracellular physical processes.

Fig. 1b illustrates the mitochondrial activation-energy ranges relevant to this argument. The correspondence is not a narrow resonance but a broad overlap of energy domains: the dominant solar photon flux (approximately 0.5–1.1 eV), characteristic biospheric activation energies, approximately 0.6–0.7 eV [80], and effective mitochondrial electron-transfer barriers all lie within the same sub-electron-volt regime. This alignment of ecological energy flux and biochemical activation energies suggests that mitochondrial electron-transfer kinetics operate within a physical environment where naturally available solar photons occupy the same energetic domain as the barriers that regulate metabolic reactions. This is not merely a coincidence of spectra, but a convergence of physical and biological constraints linking the solar radiative environment, the optical transport properties of tissue, and the energetic landscape of metabolism.

The resulting comparison shows that available solar photon power and basal metabolic demand scale across body mass in a broadly parallel fashion. Even allowing for limited photon transport efficiency and intermittent exposure, this correspondence suggests that ambient IR radiation could contribute to metabolic regulation. Because activation barriers regulate biochemical reaction rates, their characteristic energy scale also defines the range over which environmental thermal, chemical or photonic perturbations can affect cellular dynamics.

Taken together, these considerations suggest that solar photon flux, tissue optical transport, and mitochondrial electron-transfer energetics occupy overlapping physical regimes across multiple biological scales. The implications of this convergence for our understanding of photometabolism are considered below.

## 6. Discussion

### 6.1 Activation barriers as guardians of metabolic homeostasis

The mechanism developed above shows how Marcus theory [63,64] provides a quantitative description of electron-transfer reactions within a fluctuating molecular environment. It therefore offers a physically consistent framework by which environmental photons may influence metabolic kinetics without contributing directly to metabolic free energy.

Marcus theory, however, does not address a deeper biological question: why does metabolism employ activation barriers at all? An answer was foreshadowed by Schrödinger in *What is Life?* [81], where he argued that the defining characteristic of living systems is their ability to maintain an ordered, dynamic state far from thermodynamic equilibrium across timescales extending from molecular events to the lifetime of the organism. Such persistence requires not only a continuous supply of free energy but also a kinetic architecture capable of preventing spontaneous equilibration.

From this perspective, the activation barrier performs a dual function. At the molecular level, it regulates the rate of electron (and proton) transfer, as described quantitatively by Marcus theory. At the system level, it preserves the metastable organisation required for homeostasis by preventing uncontrolled relaxation towards equilibrium. Without such kinetic constraints, redox partners would rapidly equilibrate, proton gradients would dissipate, metabolic flux would become uncontrolled, and the organised state required for life could not be sustained.

Marcus theory and Schrödinger’s thermodynamic perspective may therefore be viewed as complementary descriptions of the same physical system. Marcus explains the kinetics of barrier crossing — the physical process — whereas Schrödinger explains the biological necessity for the barrier itself. In this sense, activation barriers are the kinetic guardians of metabolic homeostasis.

### 6.2 Biological architecture and the reorganisation coordinate

Marcus theory provides a general physical description of electron transfer in condensed matter, irrespective of whether the surrounding medium is a simple solvent, an electrochemical interface, or a biological macromolecule. Living systems, however, differ fundamentally from most non-biological condensed-phase environments. Over approximately four billion years of evolution, the molecular architecture surrounding each electron- and proton-transfer step has been progressively refined to optimise metabolic function. This raises a natural question: has evolution refined not only the chemistry of metabolism, but also the reorganisation coordinate that governs electron-transfer kinetics?

The reorganisation energy, λ, is a central environmental parameter in Marcus theory. It represents the free energy required to rearrange the surrounding molecular environment from its equilibrium configuration before electron transfer occurs, even when there is no net change in chemical free energy. This reorganisation includes changes in bond lengths and angles, local electric fields, hydrogen-bond networks and the orientation of surrounding water molecules, all of which contribute to the activation barrier.

In the condensed-phase systems for which Marcus theory was originally developed, these reorganisational motions generally arise from the collective response of large numbers of solvent molecules surrounding the reacting species. The fluctuating environment is therefore largely statistical, reflecting the average behaviour of many molecular degrees of freedom. In contrast, biological electron transfer occurs within highly organised molecular assemblies comprising proteins, cofactors, membrane lipids, confined water, and hydrogen-bond networks that have been progressively refined by evolution. Rather than occurring within a homogeneous solvent, each electron-transfer step in biology takes place within a precisely organised nanoscale environment whose architecture has itself become an evolutionary adaptation [82].

The extraordinary molecular organisation of biological electron- and proton-transfer systems has implications beyond efficient catalysis. If the dominant reorganisation coordinate is governed by a relatively small number of highly coordinated molecular degrees of freedom, weak perturbations of this local environment may exert a proportionately greater influence on electron-transfer kinetics than would be expected in a disordered condensed-phase system. In this view, biological organisation provides a highly refined molecular architecture within which Marcus principles operate.

This perspective suggests that evolution may have progressively optimised not only the chemical components of metabolism but also the structural environment in which metabolic electron- and proton-transfer occur. The proteins, cofactors, membrane lipids, hydrogen-bond networks, and confined water surrounding each reaction centre are therefore viewed not merely as structural components, but as integral elements of the kinetic machinery that governs metabolic flux.

Within such an organised environment, weak vibrational excitation arising from naturally available solar photons may influence the probability of barrier crossing without altering the biochemical pathways themselves. The resulting modulation would be expected to be subtle at the level of individual electron-transfer events, yet capable of producing biologically significant consequences through the continual operation of vast numbers of coupled reactions and integration within cellular regulatory networks. Such behaviour is a defining characteristic of complex biological systems and provides a plausible physical context in which the experimental observations reviewed in this paper may be understood.

### 6.3 Ecological implications

If near-IR photons contribute to metabolic regulation, then the spectral environment experienced by organisms becomes an ecological variable rather than simply an energetic background.

A striking ecological manifestation of this spectral structure is found in the forest understorey. While visible light is strongly attenuated by chlorophyll and accessory pigments, plant leaves reflect and scatter most radiation between 700 and 1300 nm to avoid thermal overload [61]. As a result, forest floors are often visibly dim but markedly enriched in near-IR, sometimes by an order of magnitude or more. This distinctive spectral environment may contribute to ecological niche differentiation and could provide a biophysical context for some documented physiological and psychological benefits of forest immersion.

Natural habitats differ enormously in their near-IR environments through canopy structure, cloud cover, vegetation, water, snow, time of day and season. These naturally occurring variations may therefore represent an underappreciated dimension of ecological adaptation and environmental physiology.

### 6.4 Beyond the BTTW

The present hypothesis has focused primarily on near-IR photons within the biological tissue transmission window (BTTW; approximately 700–1300 nm), where solar radiation penetrates deeply into living tissue [24] while overlapping the highest-flux region of the terrestrial solar photon distribution. This spectral region provides the clearest opportunity for direct interaction between environmental photons and mitochondrial architecture throughout the body. This emphasis should not be interpreted as implying that longer-wavelength IR radiation beyond approximately 1300 nm is biologically unimportant. Although absorption by liquid water increases progressively with wavelength, the atmosphere remains partially transparent across broader IR windows and the terrestrial solar photon flux remains substantial, with energy deposition becoming increasingly confined to superficial tissue layers.

Within the present framework, these photons may contribute to biology through mechanisms distinct from those proposed for the near-IR, including local effects on vascular function, immune signalling, reactive oxygen species, and other secondary pathways capable of propagating systemic physiological responses. The existence of abscopal effects following local IR irradiation suggests that biological signalling is not necessarily limited by the depth of initial photon penetration. The physiological significance of these longer-wavelength solar photons under natural conditions remains largely unexplored and represents an important direction for future investigation.

## 7. Conclusions

Throughout most of Earth’s history, the spectral distribution of sunlight has changed comparatively little, even as the luminosity of the evolving Sun increased substantially. During this time, life has evolved within a stable solar spectral environment. While biology has long recognised the importance of this environment for photosynthesis and, more recently, for vision and circadian regulation, its possible influence on the physical processes underlying metabolism has received comparatively little attention.

The evolutionary requirements of bioenergetics differ fundamentally from those of vision. Vision requires the extraction of environmental information through specialised photoreceptors, whereas metabolism is a universal property of living cells. If mitochondrial electron-transfer kinetics evolved within this persistent solar spectral environment, then that environment may represent a previously overlooked component of the physical context in which metabolism operates.

This paper has been built on the convergence of observations drawn from stellar astrophysics, atmospheric physics, tissue optics, electron-transfer theory and physiology—disciplines that have developed largely independently. Taken individually, none of these observations establishes the proposed mechanism. Taken together, however, they define a coherent physical framework in which the spectral environment, the energetics of metabolism and the optical properties of living tissue converge within the same characteristic energy domain.

Throughout its evolutionary history, terrestrial life has evolved beneath a remarkably persistent stellar spectrum. The present work suggests that this enduring spectral environment may have become more than the backdrop to evolution: it may form part of the physical context within which metabolism itself evolved and continues to operate. Whether this hypothesis proves correct in detail or requires substantial revision, we hope it encourages a broader dialogue between astrophysics, photobiology, biophysics and evolutionary biology concerning the role of the spectral environment in the physical principles that govern the organisation and evolution of life.

## Acknowledgments

RAEF and RS gratefully acknowledge the late Professor Rudolph A. Marcus for his generous encouragement, his acute interest in this project, and the insights he shared during a series of delightful video and voice-memo discussions in the winter of 2025–26, which helped shape the conceptual framework of this paper. We thank Professors Richard Weller and Andrew Gow for their valuable discussions of nitric oxide biology and its relevance to light-mediated physiological responses. Our association with the Guy Foundation provided access to the international expertise in quantum biology represented in its biannual seminar series, from which we benefited greatly.

The authors used OpenAI’s ChatGPT as an editorial and analytical assistant during the preparation of this manuscript. It was used to assist with language refinement, structural editing, literature organisation and, in particular, interdisciplinary discussion. All scientific interpretation, hypotheses, analyses and final editorial decisions were made by the authors, who take full responsibility for the content of the manuscript.

